# Wistar rats raised in an affectionate environment display lifesaving-like behaviors while distinguishing life from death

**DOI:** 10.1101/2023.08.08.552239

**Authors:** Kanta Mikami, Yuka Kigami, Tomomi Doi, Mohammed E. Choudhury, Yuki Nishikawa, Rio Takahashi, Yasuyo Wada, Honoka Kakine, Mayu Kawase, Nanae Hiyama, Hajime Yano, Naoki Abe, Toshihiro Yorozuya, Tasuku Nishihara, Junya Tanaka

## Abstract

It is generally believed that humans are the only species that values life. However, it is not well understood whether animals have a nature that values life. In this study, we attempted to determine whether male Wistar rats have this nature. Normally-reared rats did not show lifesaving-like actions towards anesthesia-induced comatose rats, although they seemed to distinguish life from death. Considering the possibility that different rearing conditions may foster a life-valuing nature, male Wistar rat pups were reared in several ways: normal rearing, loving rearing (LR; rats reared as if they were cute pets), rearing in an enriched environment, reared with gentle stroking of the back, and normal rearing of offspring of rats raised under LR conditions. When placed in an anxiety-producing environment, only LR rats escaped into the hands of the person who reared them, indicating attachment. Only the LR rats displayed lifesaving-like actions towards unknown comatose rats or drowning pups. LR rats also stopped attacks by biting ICR mice that were attacking C57BL/6 mice. Thus, rearing in an affectionate environment may foster a life-valuing nature, even in rats, suggesting that the valuing of life may be neither innate nor human-specific.

## Introduction

Even in an affluent world, war and murder persist, and continue to cause death and misery. Perhaps, underlying war and murder, there is a disorder in the formation and functioning of the “life-valuing nature.” If the processes involved in fostering the nature valuing of life could be biologically and experimentally revealed, it would be of great help in deterring undesirable of human behaviors. Therefore, it is necessary to develop animal models to evaluate the valuing of life. However, animals other than humans are not generally considered to have this nature; thus, there are no established animal models for such research.

In the present study, we investigated whether male Wistar rats have a life-valuing nature. If rats have this nature, they would try to rescue other living beings in life-threatening conditions, even if they are unknown conspecifics or of another species. We further considered that a prerequisite for a life-valuing nature is the ability to distinguish between life and death. Therefore, in this study, we first conducted a triage test (TT) to examine whether the rats could distinguish between life and death (Fig. 1A). Triage is the process of deciding which individual should be treated first based on their condition. For the TT, two Wistar littermate rats of 8–9 weeks of age were prepared that were unknown to the test rats; one was euthanized by carbon dioxide inhalation and the other was deeply anesthetized. Most of the normally reared rats spent a longer duration around the comatose rats than the euthanized rats, suggesting that the rats could distinguish between life and death, but we still could not confirm whether they valued life.

**Figure 1.**
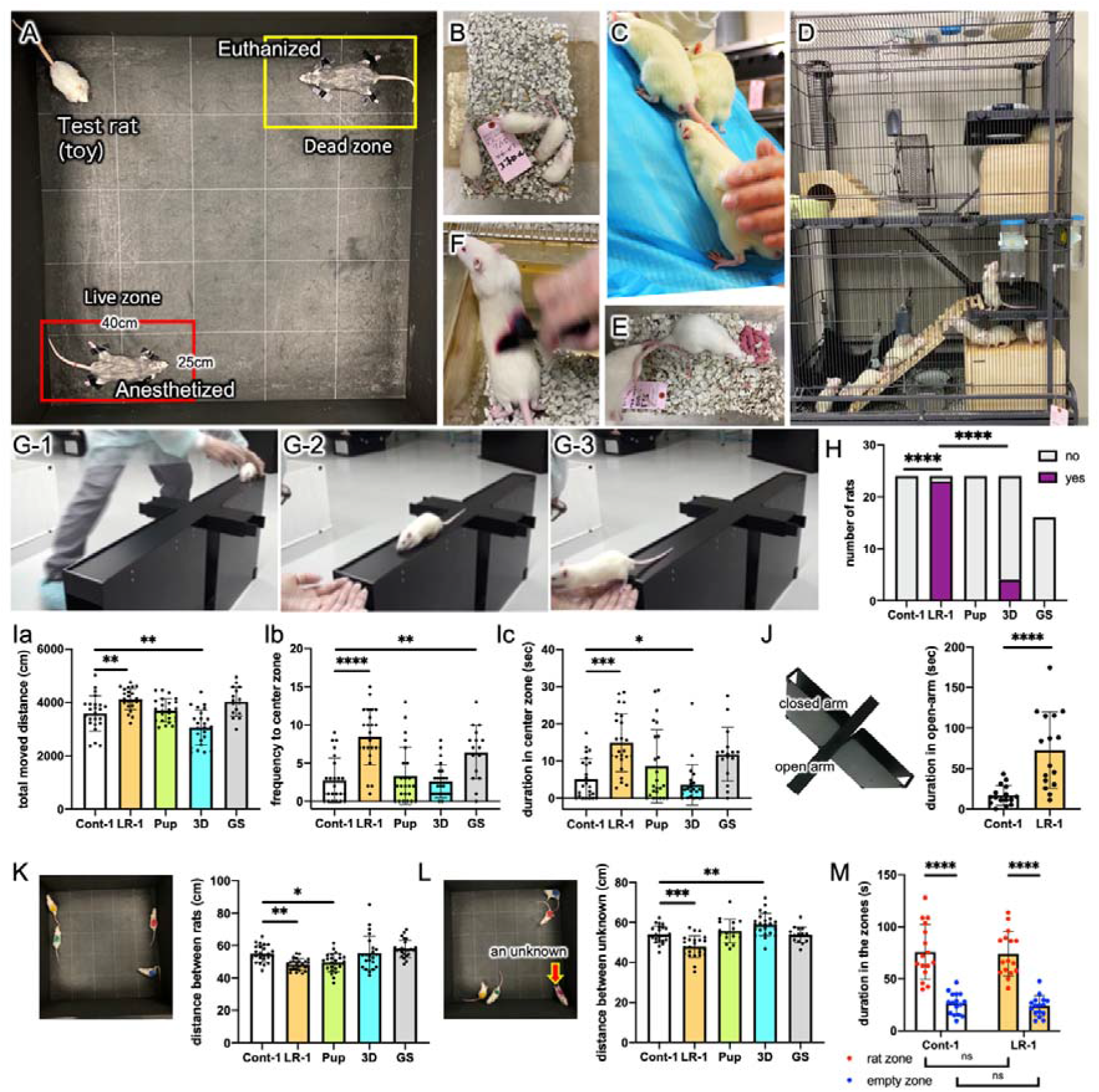
Rearing methods and the results of fundamental behavioral tests. A) TT setting. B) Rearing of Cont-1 rats nursed by their parents until PND21. Four male littermates were subsequently reared in one cage. C) LR: a caretaker playing with rats. D) 3D: group rearing of 12 rats of the same age in a large 3D cage with an enriched environment. E) Pup: pups born and raised until PND21, all reared in LR condition. F) GS: rats gently stroked on the back with a cosmetic brush. G) An LR-1 rat on AtT: the caretaker carefully placed the rat on one end of the open arm of EPM (G-1), moved to the other end, and held out both hands calling “Come!” (G-2); if the rat’s four legs were on the person’s hands within 90 s, the session was successfully over (G-3). H) AtA: almost all LR-1 rats got on the hands of the caretaker. I) OFT: total moved distance (Ia), frequency entering the center zone (Ib), and the duration staying there (Ic). J) EPM: LR-1 rats stayed longer on the open arms than Cont-1 rats. K) GOFT: four cage mates were released to the OF, and the mean distance between rats was measured for 5 min. L) GOFT with an unknown rat: mean distance between the cage mate and the unknown rat. M) 3CT: the time spent in the area around the cylinder containing an unknown rat and the empty cylinder was compared. *n* = 24, except for the GS group (*n* = 16). Values are in mm. *n* = 16 for EPM and 3CT. (H), χ^2^ test and Fisher’s exact test; (I, K, L), one-way ANOVA and Tukey’s post hoc test; (J), unpaired two-tailed *t*-test; (M), two-way ANOVA and Tukey’s post hoc test.

An unfavorable upbringing environment is clearly linked to the commission of murder and other heinous acts ^1,2^. In an animal study, rats raised in an unfavorable environment were more likely to kill mice ^3^. Mice raised in solitary environments attack other mice ^4-6^. These reports suggest that a life-valuing nature can be fostered by a favorable nurturing environment. Hence, we hypothesized that a nurturing environment could influence the life-valuing nature and behaviors of rats. To test this hypothesis, we employed a loving rearing (LR) method, in addition to normal control rearing and three other rearing methods. In the LR method, caretakers treated rats as if they were their pets, by playing and talking with them for 15 min to 1 h almost every day after weaning. The rats reared by the LR method (LR rats) displayed apparent familiarity not only with their caretaker but also with other humans. In comparison to normal control rats, LR rats more frequently contacted comatose rats in the TT. Furthermore, LR rats displayed lifesaving-like behaviors in response to drowning unknown rat pups. When LR rats experienced deadly bullying of C57BL/6JJc1 (BL6) mice by solitarily reared Jc1:ICR (ICR) mice, the LR rats stopped the bullying of the BL6 mice, which were smaller in size than ICR mice. These results suggest that a life-valuing nature is not innate, rather it was acquired when fostered by the LR method.

## Results

### Ability of male Wistar rats to distinguish life from death

To determine whether the rats could distinguish between life and death, we created a triage test (TT) (Fig. 1A). For the TT, two Wistar littermate rats (age: 8–9 weeks) that were unknown to the test rats were prepared: one was euthanized by carbon dioxide inhalation; the other was deeply anesthetized with an intraperitoneal injection of a mixture of medetomidine hydrochloride, midazolam, and butorphanol tartrate. The two rats were firmly placed on the diagonal perimeter of a 1 m^2^ open field (OF) by sticking a piece of black vinyl tape on each limb. Most of the normally reared rats spent longer around the comatose rats than the euthanized rats, suggesting that the rats could distinguish between life and death, but we still could not confirm whether they had a life-valuing nature because the rats did not show apparent lifesaving-like behaviors.

### Five rearing methods

We hypothesized that a nurturing environment can affect the nature and behavior of rats. To test this hypothesis, we employed five rearing methods:1) normal control rearing (Cont-1; Fig. 1B), 2) loving rearing (LR) by caretakers (LR-1; Fig. 1C; Fig. S1), 3) group rearing of 12 rats in a large 3D cage in an enriched environment (3D; Fig. 1D), 4) group of offspring born to LR parents (pup; Fig. 1E), and 5) gentle stroking group (GS; Fig. 1F). In particular, the rearing method used in the 3D group suppresses hyperactivity and inattentive behaviors in attention-deficit/hyperactivity disorder model rats ^7^. In the GS group, rats were gently stroked daily from the neck to the back with a cosmetic brush; this rearing method has been reported to improve social skills in mice ^8^. All rats were self-bred; mature male and female Wistar rats were cohabited, and pups were nursed with their parents until postnatal day (PND) 21. For four LR-1 rats in a cage, two caretakers (a male professor in his 60s and a female undergraduate in her 20s) cared for and played with the rats as if they were their pets for 30 min to 1 h 6 or 7 times per week. LR was conducted from PND21 to the end of the behavioral experiments. The caretakers were not present during the behavioral experiments, except during the newly created attachment test (AtT). The AtT partly resembles the human approach test that evaluates the attraction of rats to a human hand, which has been reported elsewhere ^9^. However, the AtT was performed with only an open arm (100 × 10 × 50 cm) of an elevated plus maze (EPM) apparatus in an unknown laboratory, triggering a sense of anxiety under bright illumination (1,220 lx) to examine whether they escaped into the caretaker’s hands (Fig. 1G). In the AtT, the male caretaker initially placed the rats on one end of the arm, moved to another end, and called the rat, “Come here!” Of the 24 rats, no Cont-1 rats, 23 LR-1 rats, no pups, and four 3D rats climbed onto the hands of the caretaker within 90 s. In contrast, none of the 16 GS rats exhibited this behavior. Clearly, the LR-1 rats were exceptionally attached to the caretaker but also appeared to feel a sense of familiarity with two unknown behavioral experimenters (male and female undergraduates in their 20s; Fig. S2). In contrast, Cont-1 rats did not show any familiarity with the caretaker who took care of them each day (Fig. S2).

### Behavioral characterization of rats reared in the different methods

LR-1 rats were more active and less anxious, while 3D rats were less active and more anxious than Cont-1 rats according to the OF test (OFT) and the EPM results (Fig. 1I and 1 J) ^10^. Moreover, four rats from the same cage underwent grouped OFT (GOFT) (Fig. 1K); the mean distance between the two rats was the shortest in the LR-1 group. When an unknown male Wistar rat of the same age was added to these four rats, the mean distance to this unknown rat was also the shortest in the LR-1 group (Fig. 1L). The three-chamber test (3CT) showed no significant difference between the Cont-1 and LR-1 groups (Fig. 1M) ^11^. Taken together, LR-1 rats were relatively more social than Cont-1 rats.

### The triage test

In the TT, a live zone was set around the comatose rats and a dead zone was set around euthanized rats; the time spent in each zone was measured (Fig. 1A, 2A). After the test rats were released from another corner, their behavior was recorded for 5 min. The total distance moved decreased only in the 3D group (Fig. 2B). The Cont-1, LR-1, pup, and GS groups stayed longer in the live zone than in the dead zone, whereas only the 3D group stayed longer in the dead zone (Fig. 2C, 2D). We manually counted the number of contacts made by the forepaw, nose, or mouth of the test rats to the head and body of the euthanized or comatose rats (Fig. 2E, 2F, 2G; Fig. S3). LR-1 rats contacted the head of comatose rats and the head and body of euthanized rats more frequently than Cont-1 rats (Fig. 2G). In addition, the heads of comatose rats were contacted more frequently by LR-1 rats than by euthanized rats (Fig. 2H). The time spent in the OFT center zone (an index of anxiety) was not correlated with the number of contacts of the heads of comatose (Fig. 2Ia) and euthanized rats (Fig. 2Ib) by the LR-1 rats.

**Figure 2.**
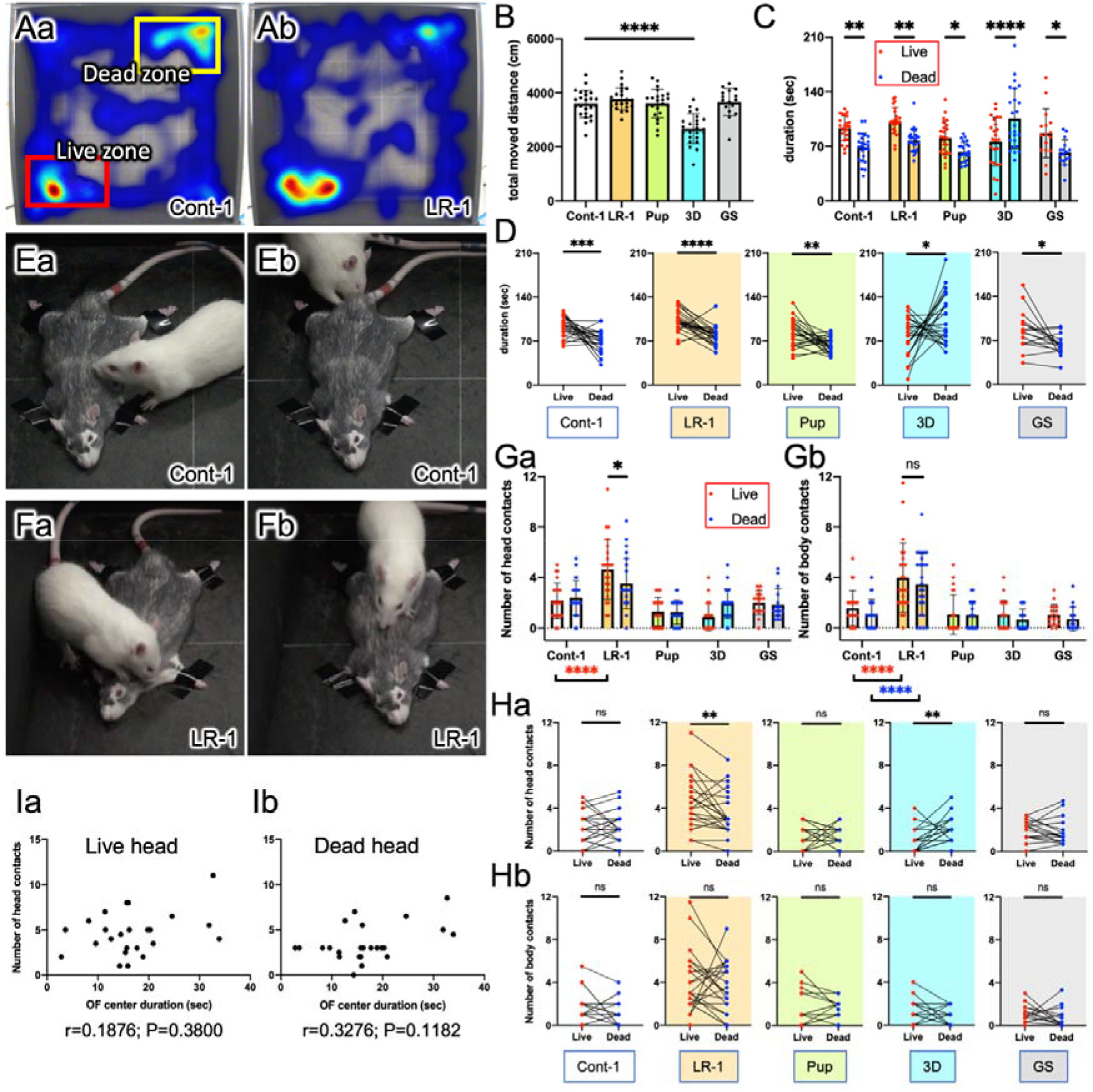
Triage test. A) Setting of live and dead zones in the OF arena and the representative heat maps showing the duration of staying in the OF by a Cont-1 rat and a LR-1 rat. B) Total moved distance in the TT. C) Duration of staying in the live or dead zones. D) Comparison of the duration of staying in the live and dead zones by individual rats. E, F) Captured images from video recordings around a comatose rat accompanied by a Cont-1 rat (E) or a LR-1 rat (F) whose total duration in the live zone was approximately 2 min. Only the LR-1 rat frequently contacted the head (Fa) or body (Fb) of the comatose rat. G) Number of test rats’ contacts with the heads (Ga) and bodies (Gb) of the comatose and euthanized rats. H) Comparison of the number of test rats’ contacts with the heads (Ha) and bodies (Hb) of the comatose and euthanized rats. I) Correlation between the duration in the center zones in the OFT and the number of contacts with the heads of the comatose (Ia) and euthanized (Ib) rats by LR-1 rats. *n* = 24, except for GS group (*n* = 16). (B), one-way ANOVA and Tukey’s post hoc test; (C, G), two-way ANOVA and Tukey’s post hoc test; (D, H), paired two-tailed *t*-test; (I), Spearman correlation analysis.

The 16 LR-2 rats shown in Figs. 3 and 4 were reared with love only by the male caretaker for 15 min per four rats in a cage almost every day. The LR methods were the same as those for the LR-1 group, but all LR-2 rats were called “Rick,” the name of the caretaker’s dog. When called “Rick” in the AtT, one of the 16 rats in the Cont-2 group and all 16 rats in the LR-2 group got onto the hands of the caretaker (Fig. 3A). In the OFT, LR-2 rats moved shorter distances than LR-1 and Cont-2 rats, and the LR-2 group did not significantly differ from the Cont-1 and Cont-2 groups with regard to the frequency of center zone entries and the time spent in the center zone, which were significantly lower and shorter, respectively, in comparison to the LR-1 group (Fig. 3B). The duration of stay in the open arm in the LR-2 group was significantly shorter than that in the LR-1 group, with no significant difference between the Cont-1 and Cont-2 groups (Fig. 3C). Thus, LR-2 rats differed slightly from LR-1 rats in terms of activity and anxiety.

**Figure 3.**
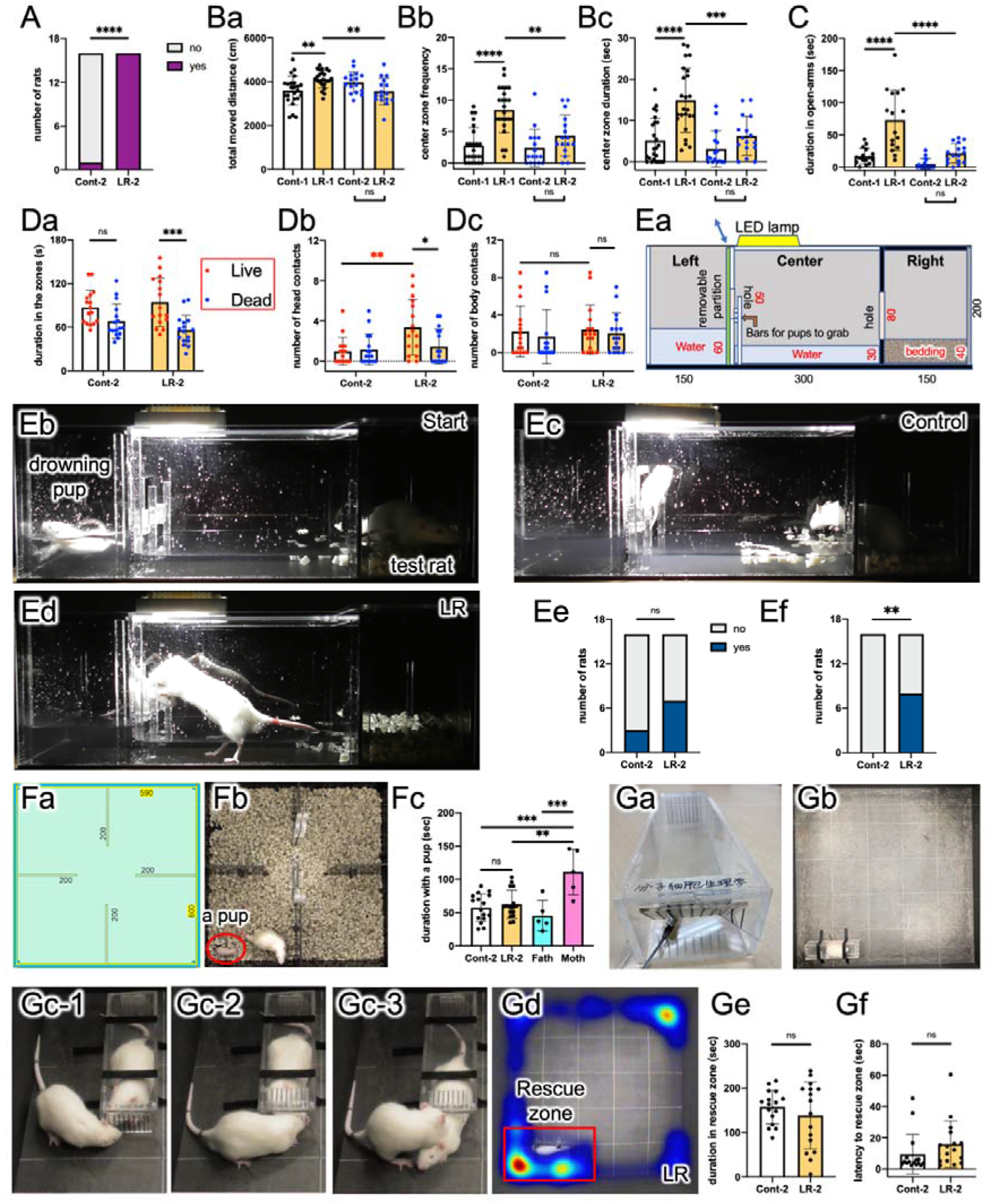
Cont-2 and LR-2 rats. A) AtT. B) OFT: total distance (Ba), duration in the center zone (Bb) and frequency entering the zone (Bc). C) EPM. Cont-1 and LR-1 data in (B) and (C) are the same as shown in Fig. 2. D) TT: duration in live zone (Da) and number of contacts to the head (Db) and the body (Dc). E) DT: the apparatus (Ea). Values are in mm. The pup and the test rat (Eb). Response of a Cont-2 rat (Ec) and an LR-2 rat (Ed). The number of rats touched the boundary during habituation (Ee) and those contacted the drowning pups (Ef). F) 4RT: the 4RT box (Fa). Colocalization of a test rat and a pup (Fb). Duration in the same room (Fc). G) RarT: the apparatus (Ga) and that on the OF arena (Gb). A rescued rat (Gc-1 to Gc-3; fig. S7). Rescue zone and a representative heatmap showing 5 min movements of an LR-2 rat (Gd). Rescue zone duration (Ge) and the frequency entering the zone (Gf). *n* = 16. (A, E), χ^2^ test and Fisher’s exact test; one-way (B, C, F) or two-way (D) ANOVA and Tukey’s post hoc test; (G), unpaired two-tailed *t*-test.

**Figure 4.**
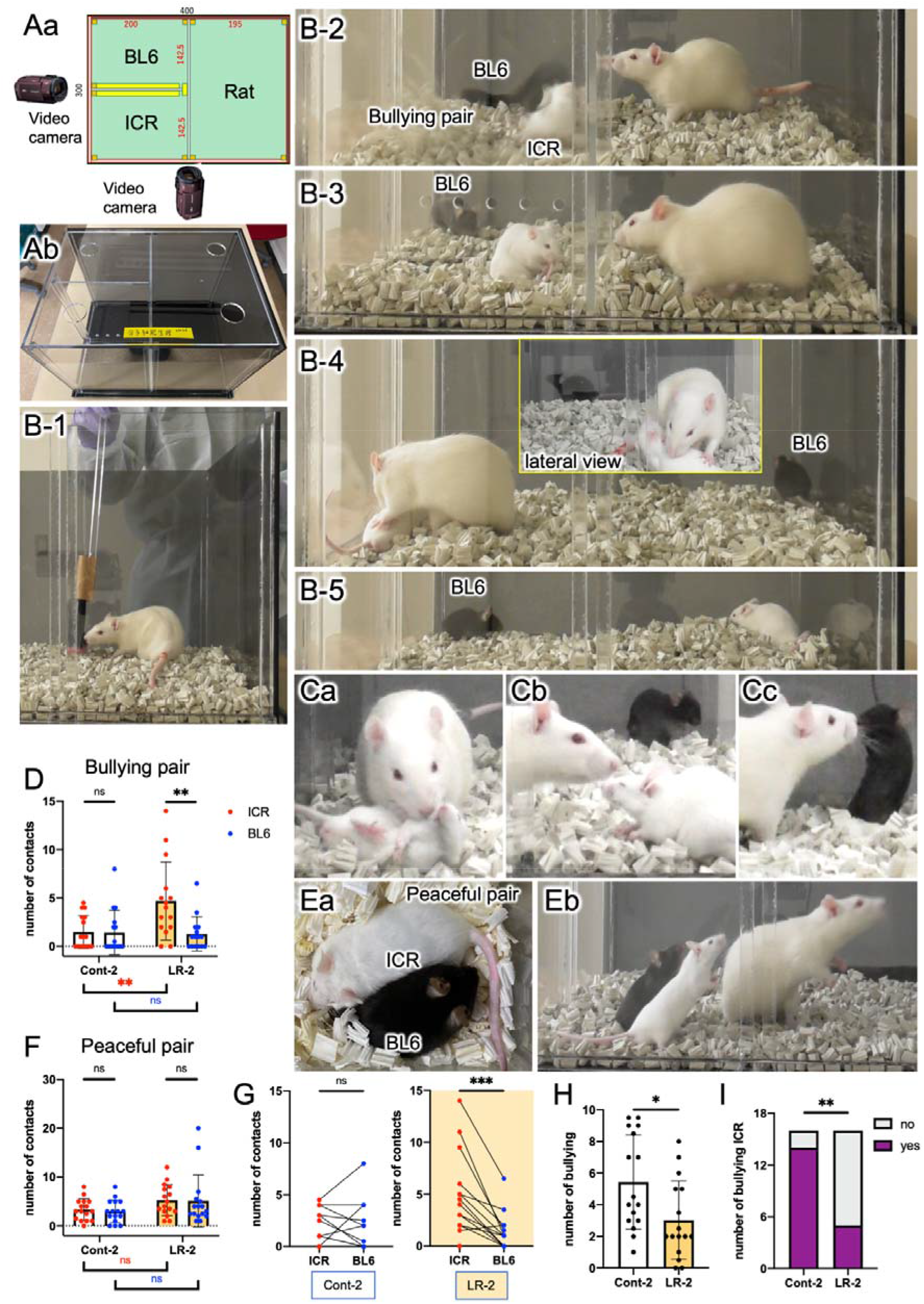
Bullying test (BT). A) BT apparatus. Values are in mm. B) The observation procedure in the case of a bullying pair. C) Behavior of another animal group. The LR-2 rat appeared to admonish the ICR to stop bullying (Ca, Cb). BL6 was distancing itself from both. The LR-2 rat smelled but did not frequently contact BL6 (Cc). D) The number of times the rats contacted ICR or BL6 with their forepaws. E) In the peaceful pair, there were no fights at all in the absence (Ea) or presence (Eb) of rats. F) The number of contacts in the peaceful pairs did not significantly differ between the LR-2 and Cont-2 groups. G) The LR-2 rats contacted ICR of the bullying pair more frequently than the Cont-2 rats. H) The ICR reduced the number of attacks in the presence of the LR-2 rats. I) The number of the ICR mice attacked the BL6 within 1 min after rat removal. Cont, *n* = 16; LR, *n* = 16; only the LR-2 group data in D, *n* = 15. (D, F), two-way ANOVA and Tukey’s post hoc test; paired (G) or unpaired (H) two-tailed *t*-test; (I), χ^2^ and Fisher’s exact test.

In the TT, LR-2, but not Cont-2, rats stayed longer in the live zone than in the dead zone (Fig. 3Da, Fig. S4). LR-2 rats contacted the heads of comatose rats more frequently in comparison to Cont-2 rats (Fig. 3Db). LR-2 rats contacted the heads of comatose rats more frequently than they contacted the heads of euthanized rats. A statistical analysis of the TT results showed that the behavior of the LR-1 and LR-2 groups was similar.

### The drowning test

To directly observe lifesaving-like behaviors, we developed a drowning test (DT) using a homemade acrylic box with three chambers (Fig. 3Ea) in which an unknown pup (PND16) was drawn. We filled the left chamber with water to a depth of 6 cm and the center chamber (30 cm wide) to a depth of 3 cm (Fig. 3Ea). An LED light was placed above the center chamber with an illuminance of 11,100 lx directly below it. The right chamber was separated from the center chamber by a black opaque acrylic panel and covered with a 4-cm-deep floor bed. The rear and right outer sides were surrounded by grey translucent acrylic panels. Experimenters observed rats behind these translucent walls. To allow the rats to enter and exit, we drilled a square hole between the left and right chambers and the central chamber. Furthermore, a transparent acrylic partition can be inserted into the left chamber to close the entrance to the center chamber. If the test rat tried to rescue the drowning pup, it needed to overcome the aversive obstacles of the water and bright illumination (Fig. 3E, Fig. S5).

In the DT, the test rats were placed in the right chamber for 3 min of habituation. Although no significant difference was noted, a greater number of LR-2 rats entered the center chamber in comparison to the Cont-2 rats (Fig. 3Ee). After being placed in the left room (with a water depth of 6 cm), the unknown PND16 pup moved violently in an attempt to escape (Fig. 3Eb), and the test rats could observe this situation for 1 min. Following the removal of the partition, half of the LR-2 rats touched the drowning pup within 30 s; however, none of the Cont-2 rats approached (Fig. 3Ec, 3Ed, 3Ef). Mother rats were able to rescue their pups by placing them in their mouths (Fig. S6), whereas male rats were unable to do this; thus, LR-2 rats only snuggled up to the drowned pups (Fig. S5).

To examine whether the rescue-like behaviors shown during the DT emerged out of concern for their lives in the unknown pups or out of interest in them, we established a 4-room test (4RT) using a homemade acrylic box, which had an area of approximately 60 cm^2^ with 45-cm-high walls. The interior was divided into four compartments using opaque black acrylic panels (Fig. 3Fa and 3Fb), and the bottom was covered with a floor bed. We video-recorded the movements of the test rat and pup for 5 min and measured their colocalization duration in the same room. For comparison, we also measured the duration of the colocalization of either the mother or the father rats with their pups (Fig. 3Fc). The 4RT results showed that the LR-2 rats did not show any particular interest in the pups; possibly, they judged that the situation did not require immediate life-saving actions.

### Test to evaluate the empathy of the rats

To investigate whether the behavior of LR rats toward unknown comatose rats and drowned pups was related to empathy, we observed the behavior of LR-2 and Cont-2 rats toward an unknown restrained male rat of the same age (Fig. 3G). This was called the rescuing a restrained rat test (RarT). This is similar to a previously established experiment ^12^; however, the restrained rectangular container was of simpler construction, with front and rear doors that could be easily opened by pushing from the outside (Fig. 3Ga). However, if the restrained rat attempts to escape from the inside, the door will not open, and cooperation between the two rats is required. The container was placed in the same position as the live zone in the TT. The movements of the test rats were recorded using a video tracking system and video camera (Fig. 3Gb). Within 5 min, none of the test rats rescued the restrained rat; notably, one Cont-2 rat successfully rescued the restrained rat within 6 min (Fig. 3Gc; Fig. S7). To evaluate empathy leading to rescue behavior, we measured the duration that the test rats spent in the rescue zones, which is the same zone as the live zone in the TT, and their frequency of entry into the rescue zones for the first 5 min. However, these data did not differ to a statistically significant extent between the Cont-2 and LR-2 groups (Fig. 3Gd, 3Ge, and 3Gf). Considering that rescue behavior in the RarT is based on the rats’ empathy ^12,13^, the LR method did not appear to strengthen their empathy.

### The bullying test

Jc1:ICR (ICR) mice raised in an isolated environment often attack C57BL/6JJc1 (BL6) mice by biting, sometimes leading to severe injury or death. We considered that the attacks by ICR mice on BL6 mice amounted to deadly bullying, and we observed rats’ behavior toward bullying in the newly created bullying test (BT; Fig. 4; Fig. S8 and S9). The BT apparatus included three rooms that could be partitioned by acrylic transparent removable boards with a row of holes (diameter: 1 cm) drilled 5 cm from the bottom with 2-cm-thick bedding (Fig. 4A). These rooms were designated as ICR, BL6, and Rat rooms, respectively. In the experiment, we first selected an ICR mouse with a high frequency of attacks using this apparatus. Peaceful pairs (no aggression between mice) were also prepared. Video cameras were placed at the front and sides of the apparatus. During the test, the test rat was first placed in an empty apparatus and allowed to move freely for 2 min while being habituated to the place and a cosmetic brush, which was attached to the end of a 40-cm-long acrylic rod (Fig. 4B-1, Fig. S8 and S9). This brush was used to separate animals in the case of a severe attack by a mouse or rat. The bullying pair was then placed in the mouse zone with the central partition in place, and the test rat observed the attacks by the ICR mouse on the BL6 mouse through the partition for 3 min or until the ICR mouse made five attacks (Fig. 4B-2). Considering that the rats may have become agitated and aggressive by watching the bullying, we placed a partition between the mouse rooms to stop the attacks and waited for 2 min for the agitation of the rats to subside (Fig. 4B-3).

Then, we removed all partitions to allow the three animals to make contact with each other and observed them for 3 min (Fig. 4B-4). After the rat was removed, we assessed whether the ICR mouse attacked within 1 min. From these video recordings, we measured the number of times the test rats contacted the ICR mouse (Fig. 4B-4, 4Ca) or the BL6 mouse with their forepaws, and the number of times the ICR mouse attacked the BL6 mouse in the presence of the rats. One LR-2 rat that lunged at the ICR mouse promptly after the removal of the partitions was frequently stopped by the brush; this result was excluded from the bullying pair data (Fig. 4D; Fig. S9), because of the difficulty in counting the number of contacts. In contrast, Cont-2 rats did not actively contact bullying ICR mice.

In the bullying pair, LR-2 rats frequently contacted the ICR mouse, but the number of contacts with BL6 mouse did not differ from that of the Cont-2 rats (Fig. 4D). Some LR-2 rats appeared to admonish the ICR mouse not to do the bully (Fig. 4Ca, 4Cb; Fig. S8) and made less direct contact with the BL6 mouse (Fig. 4 Cc). Moreover, LR-2 rats significantly reduced the number of attacks by ICR mice during their stay (Fig. 4H) and after their removal (Fig. 4I). In the case of peaceful pairs, the number of contacts with ICR and BL6 mice by both Cont-2 and LR-2 rats (Fig. 4F) did not differ to a statistically significant extent, and the animals remained calm (Fig. 4Ea).

## Discussion

TT showed that rats that had never seen death could distinguish between life and death, a probable prerequisite for a life-valuing nature. The value of life should be absolute; hence, experiments evaluating rat behaviors were performed using rats and mice that the test rats had never encountered. The apparent enthusiastic attitudes of LR rats toward lifesaving-like behaviors in TT, DT, and BT were similar to those of humans encountering the same situations, suggesting that LR rats probably have a life-valuing nature.

LR rats showed strong familiarity with caretakers, which could be recognized as their attachment to the caretaker, as shown by the behaviors in the AtT. Other rats reared in different ways did not display significant attachment. Even the GS rats showed no attachment, despite the frequent contacts with caretakers who gently stroked the rats’ backs with a cosmetic brush almost every day. The GS results suggest that physical contact alone does not produce lifesaving-like behavior. The conditions of 3D rearing may partly resemble a wild environment, because rats were free to move around, sleep where they wanted, feed where they wanted, and interacted with many conspecifics. However, only rats reared in 3D conditions stayed longer in the dead zone than in the live zone in the TT. Euthanized rats might have appeared to be safer for 3D rats than comatose rats. The results in 3D rats may indicate that wild animals do not have a life-valuing nature. The results from the pup group suggest that love from not only parents but also other surrounding individuals and the attachment of pups to their parents and surrounding individuals may be essential to foster a nature that supports the performance of lifesaving-like behaviors.

Handling of rats by humans during infancy and the post-weaning period has often been reported to reduce anxiety in rats ^14,15^. Maternal care with frequent licking and grooming decreased anxiety ^16^ and the glucocorticoid response^17^. Therefore, the prolonged handling after weaning (LR) in this study may have reduced anxiety behaviors, as shown in Figure 1. Reduced anxiety may be correlated with the lifesaving-like behaviors of the LR rats, as assessed by frequent contact with unknown comatose rats in the TT, unknown drowning pups while overcoming the obstacles of the water pool and dazzling light, and bullying ICR mice. LR-1 rats showed markedly reduced anxiety and increased contact with unknown comatose rats. However, the number of contacts was not significantly correlated with anxious behaviors, such as a reduced duration in the center zone in the OFT. Although LR-2 rats did not differ from Cont-2 rats in terms of anxiety, their lifesaving-like behaviors were apparent. These observations suggest that lifesaving-like behaviors are not the result of reduced anxiety. In addition, the handling of rats is generally said to reduce anxiety, rearing of animals in an enriched environment is reported to increase anxiety ^18^. Thus, the difference in handling methods between the LR-1 and LR-2 rats may have caused the difference in anxious behaviors.

Next, we investigated whether lifesaving-like behaviors were correlated with empathy. Empathy has been suggested to underlie the rat behavior of rescuing cage mates who cannot escape from restrained situations ^12,13^. However, in our study, the behavior of trying to rescue restrained unknown rats in the RarT showed no difference between the Cont-2 and LR-2 rats. Sociality may also be involved in increased contact with unknown animals. However, the GOFT showed that LR-1 rats socialized relatively better than Cont-1 rats, but no difference was noted in 3CT. The interest in isolated immature pups did not differ between the Cont-2 and LR-2 rats at 4RT. In the RarT, if a freely moving rat tries to rescue a restrained rat, the former is required to cooperate with the latter. Therefore, the RarT may also be used to evaluate the sociality of rats. Taken together, these results suggest that LR did not significantly alter sociality. All these considerations suggest that lifesaving-like behaviors are independent of anxiety, empathy, and sociality. From a human perspective, the isolated immature pups in 4RT and the restrained mature rats are in a critical situation, but the LR rats may have judged that the situation did not require immediate lifesaving actions.

During the observation phase of the BT, the test rats watched the biting behavior of ICR mice on BL6 mice. Shortly after removing the partition that divided the test rat and the mice, the majority of LR rats rushed to the ICR mouse. If the LR rat behaviors were born out of a life-valuing nature alone, it might have approached the BL6 mouse to protect it from the ICR mouse. Therefore, the behaviors of LR rats in the BT might involve two psychological factors: the life-valuing nature and a sense of justice that discourages the strong and helps the weak. Alternatively, a life-valuing nature may be a component of building a sense of justice.

Eight-month-old infants reportedly prefer to attack a bully in animations, indicating that antipathy toward antisocial behavior is innate ^19^. Antipathy may be correlated with a sense of justice in helping the weak. However, the present results suggest that it is likely to be acquired, including a sense of valuing life. Even 8-month-old infants should have already received love from their parents and other surroundings. In the BT situation, there was no possibility of rats being attacked by mice. In this sense, the BT may be a model for third-party punishment in rats. Third-party punishment is said to be unique to humans ^19-21^, but this notion might not be true, based on the present BT results. The nature of third-party punishment may also be acquired and nurtured by the love provided by their rearing environments.

The present results suggest that a life-valuing nature was not present in rats reared in the absence of love. LR is necessary to foster a life-valuing nature as well as attachment to caretakers in rats. LR takes a very long time, and the rats were deeply attached only to a few people who tried to rear them with love in our experience. Therefore, the LR model may not be easy to establish. However, the present LR model provides a novel method that can be used to elucidate the neuroscientific background of a life-valuing nature. This would be of great help in deterring war, murder, and other behaviors that emerge when life is not valued.

## Materials and Methods

### Animals

Breeding of rats and mice and all animal experiments conformed to the Guidelines of the Ethics Committee for Animal Experimentation of Ehime University in Matsuyama, Japan (approval No.: 05U43-2; 05U47-2; 05U46-1,2). All routine care measures, such as bedding changing, feeding, and breeding, were performed very carefully only by a male caretaker (a professor in his 60s) to avoid variation in the routine rearing of animals.

#### Rats

Two pairs of Wistar rats were purchased from CLEA Japan, Inc. (Tokyo, Japan) in 1998 and home-bred thereafter. Nine-week-old male and female Wistar rats were allowed to cohabitate, gestate, and give birth. Only males were used in our experiments. Pups subjected to our behavioral experiments were kept with their parents until postnatal day (PND) 21. After separated from their parents on PND21, pups were reared in one of the following five rearing methods: control (Cont) group, loving rearing (LR) group, pup group, rearing in a large 3D cage (3D) group, and gentle stroking (GS) group. Except for the 3D group, the pups were reared in a standard plastic cage (40 × 25 × 14 cm), with four pups per cage, and behavioral experiments were conducted when they were 8–10 weeks old. Food and water were provided ad libitum, and room lighting (240 lux near the cage) was switched on from 7:00 to 19:00, and off at other times. The temperature of the room was 25 ± 1 °C, and the humidity was approximately 55%.

#### Five rearing methods for rats

1. Cont-1 and Cont-2 groups: Even-numbered male siblings of the same litter were divided into two groups and assigned to the Cont and LR groups. The Cont group was reared under normal rearing conditions.
2. LR-1 and LR-2 groups: The LR groups consisted of a litter the same as that of the Cont group. The floor bed material made of paper was changed thrice a week, and pellet food was spread on the floor daily so that the rats could eat the food with hand. The rats were not dangled by their tails, even when changing cages. LR was provided so that the rats could maximally enjoy their life; some of the LR actions were as follows: frequently talking using words such as “cute,” “smart,” “come here,” and “see you tomorrow” as much as possible to the rats; holding and petting them, allowing them to freely play on the LR persons’ chest and belly, and allowing rats to play with other LR rats of different ages (Fig. 1C; Supplementary Fig. 1). In the LR-1 group, the rats were mainly handled by the male LR person, but occasionally by a female person (an undergraduate in her 20s) for 30–60 min per four rats in one cage almost every day. In the LR-2 group, only the male LR person performed LR for 15 min in the four rats in a cage daily. All LR-2 rats were named “Rick” and called “Rick” daily (“Rick” was the name of the LR person’s dog). The LR was started at PND21 and done between 16:30–19:00 daily until the end of the behavioral testing.
3. Pup group: Four male and four female Wistar rats, born to different parents, were reared from PND21 to PND63 using the LR method. At 8–9 weeks old, they were mated to prepare four pairs and allowed to give birth. The male pups (24 pups in total) delivered were separated from their parents on PND21, and they were then reared in the same manner as the Cont group.
4. GS group: Starting at PND21, rats’ backs were gently stroked with a cosmetic brush for 10 min daily according to a previously described method ^8^. Otherwise, the rats were reared in the same manner as the Cont group.
5. 3D group: Twelve pups were reared in a large, tall cage (95 × 55 × 115 cm) equipped with a running wheel, four stages, three wooden pens, a wooden ladder, plastic slopes, a climbing steel tube, three trays with food pellets, and two trays filled with water from PND21 until PND70 ^7^.

#### Breeding of unknown rats for behavioral experiments

The male Wistar rats subjected to the triage test (TT), Rescuing a restrained rat test (RarT), grouped open-field test (GOFT), and 3-chamber test (3CT) were born to parents with whom the test rats had never been in contact, and were subsequently separated from their parents at PND23. Thereafter, they (four animals per cage) were reared in the same manner as the Cont group and used for the behavioral tests at 8–9 weeks of age. For 4-room test (4RT) and drowning test (DT), pups born to parents that were never in contact with the test rats, were used at PND11 and PND16, respectively; their sex was not taken into account.

#### Mice

Male Jc1:ICR mice (ICR) at 9 weeks old and male C57BL/6JJc1 mice (BL6) at 8 weeks old were purchased from CLEA Japan. After the purchased ICR mice were kept for a week, with four mice per cage, they were solitary kept in a cage. The BL6 mice were kept until the end of the behavioral tests, with four mice per cage. After 7 days of rearing from the day of purchase, the BL6 mice were individually and randomly placed into the ICR cage. We then observed whether the ICR attacked the BL6 or not. Repeating this temporary cohabitation, which triggers ICR attacks, we identified 10 pairs of attacking ICR and attacked BL6. Prior to the bullying test (BT), the ICR and BL6 were also placed in the BT apparatus to confirm that the ICR attacked the BL6 within 1 min. When the ICR gave excessively violent attacks on BL6, the two animals were separated using a cosmetic brush that was attached to an acrylic rod (Fig. 4 B, fig. S9). Furthermore, two ICR-BL6 pairs that did not exhibit bullying were designated as peaceful pairs. While lighting on/off, feeding, water supply, and other conditions were the same as for the rats, the rooms for the rats and mice differed. Routine care was carefully performed only by the male LR person. The ICR and BL6 mice were used for BT until they were 13 and 12 weeks old, respectively.

### Behavioral Tests

Except for AtT, behavioral experiments were not performed by the LR persons but by other experimenters (male and female undergraduates in their 20s; fig. S2) who had not touched the animals until the behavioral tests. No handling was used to calm the rats or mice before any of the behavioral experiments, except for the LR rats. Behavioral experiments were conducted from 17:00 to 21:00. In the Cont-1, LR-1, 3D, pup, and GS groups, behavioral tests were conducted in the following order: AtT, open-field test (OFT), TT, GOFT, elevated plus maze (EPM), and 3CT. EPM and 3CT were conducted only in the Cont-1 and LR-1 groups. In the Cont-2 and LR-2 groups, the tests were performed in the following order: AtT, OFT, DT, TT, 4RT, BT, and RarT. Only one test was done in a day, with up to four tests per week. The LR person told the LR rats “go for it” before the test and “good job” thereafter, while patting the body, with an aim to maintain trustworthiness. When subjected to behavioral tests except for AtT, all rats were held by the tail.

#### AtT

AtT was conducted to evaluate whether the rats trusted the male LR person. The LR person placed a rat at one end of an open arm of the EPM (100 × 10 × 50 cm) in a brightly lit laboratory (1,220 lux on the open arm) that the rats had never been in before. Then, the LR person immediately moved to the other end of the arm and said “Come” to the rats. In the case of the Cont-2 and LR-2 groups, he called the rats “Rick.” Normally, when rats were placed at the end of the arm, the rats were held by the body, but in the case of the Cont and other groups that did not like to be touched, the rats were held by the tail. If the rats’ four legs got on the hands of the LR person within 90 s, the test was judged as successful (Fig. 1G). If the rat did not come to the hand by 90 s, we terminated the test. Some of the tests were recorded using a video camera (HC-W590MS; Panasonic, Osaka, Japan).

#### OFT

A 5 min OFT was conducted using a square box (100 × 100 cm) with 50 cm-high walls. We used a video-tracking system (Ethovision XT 14; Noldus Info. Tech., Wageningen, The Netherlands) to monitor the rats’ movement and measure the following parameters: total moved distance, mobile duration, frequency entering the center zone (60 × 60 cm), and latency to the first entry into the center zone. The illumination at the center of the OF arena was 54 lux ^22^.

#### EPM

The open and closed arms of the EPM apparatus made of black opaque acrylic boards (50 × 10 × 50 cm) were used in this experiment. Closed arms were walled in with 50 cm-high black opaque acrylic plates. The test was started by placing the rats at the center of the apparatus. The video-tracking system was used to record rats’ movements for 5 min and measure the duration of staying in the open arms. The illuminance was 53, 75, and 6.6 lux at the center of the apparatus, end of the open arms, and end of the closed arms, respectively ^10^.

#### GOFT

Four rats in the same cage or five rats in the group with four rats plus one unknown male rat of the same age were simultaneously released into the abovementioned OF (1 m square; Fig. 1K, 1L), and their behavior was recorded using the video-tracking system for 5 min. Red, blue, green, yellow, and pink were painted on the rats’ backs for recognition by the video-tracking system. The averaged distance between each rat from the four-rat group or between the unknown rat and the four rats in the case of the five-rat experiment was measured.

#### 3CT

At two opposite corners of an acrylic box (86 × 59.5 × 40 cm) as shown in the below, we placed 17 cm-diameter clear acrylic cylinders with numerous holes in the wall. One cylinder contained an unknown rat of the same age as the test rat, while the other cylinder was left empty. The test rat was then placed in the central zone partitioned by acrylic boards with a 10 × 10 cm square passageway, and their behaviors were recorded using the video-tracking system for 5 min. The rat zone was the area within 10 cm around the cylinders in which the unknown rat was placed, and the empty zone as the 10 cm around the empty cylinder. We next measured the test rat’s duration of staying in each zone. The illumination at the center of the apparatus was 52 lux.

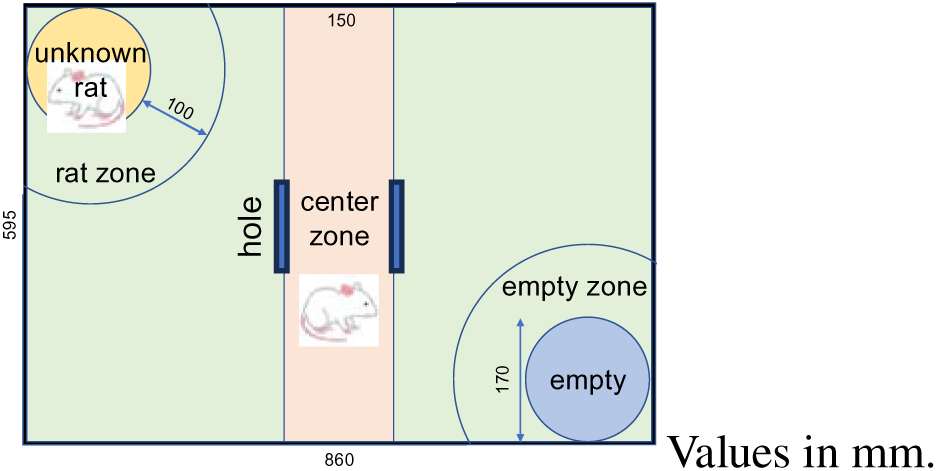

#### TT

Two littermates (male Wistar rats at 8–9 weeks old) that were unknown to the test rats were prepared per eight test rats; one was euthanized by carbon dioxide inhalation, and the other was injected intraperitoneally with 0.3 ml per 100 g body weight of a mixture of medetomidine hydrochloride (0.75 ml; Kyoritsu Pharmaceutical, Tokyo, Japan), midazolam (2 ml; Astellas, Osaka, Japan), butorphanol tartrate (2.5 ml; Meiji Seika Pharma, Tokyo, Japan), and saline solution (7.25 ml) to be comatose. The whole bodies of the two rats were painted black to prevent misidentification by the video-tracking system. These two rats were placed in the corner symmetrical to the OF, and their limbs were fixed to the bottom surface with a black vinyl tape. The illumination was the same as that in the OFT. A 40 × 25 cm area around the comatose and euthanized rats was designated as the Live zone and Dead zone, respectively (Fig. 2A). The test rats were released from the other corner (Fig. 1A), and their behavior was recorded using the video-tracking system; additionally, two video cameras were simultaneously used to record the test rats’ behavior around the Live and Dead zones. The video-tracking system was used to measure the total distance traveled and the duration of stay in the two zones. Two undergraduate students and a laboratory secretary, who were all not involved in the experiment and were unaware of the experimental conditions, observed the video recordings and measured the number of times the test rat contacted the head (above the shoulder) or body (below the shoulder) of the comatose or euthanized rat by the nose, mouth, or forepaw. From the measurements of three persons, two data with close values were extracted and averaged to obtain the results.

#### DT

An acrylic box (60 × 20 × 20 cm) with three chambers was made (Fig. 3E). The left and right chambers were 15 cm wide, and the center chamber was 30 cm wide. The rear and right sides are separated by translucent gray acrylic panels, through which an experimenter observed the animal behavior. Moreover, the center and right chambers were separated by an opaque black acrylic panel with 8 cm square hole, where the rat could pass. We also partitioned the center and left chambers by using a removable transparent acrylic panel with 5 × 7 cm rectangular hole. The left chamber was filled with water at 6 cm deep, and the center chamber at 3 cm deep. The water temperature was 23 °C–25 °C. The right chamber was covered with a dry, unused floor mat made of paper. A 15W LED lamp was placed above the center chamber. The illuminance of the box was 11,100, 6,660, 15.3, and 1,350 lux at the center chamber under the light, the left chamber, the right chamber in shadow, and the right chamber in light near the square hole, respectively. A video camera was set in front of the apparatus to record the animals’ behavior. In the experiment, rats were first placed in the right chamber and allowed to freely move in the apparatus for 3 min for habituation. After inserting a partition between the left and the center chambers, we placed an unknown rat pup of PND16 into the water in the left chamber and left it there for 1 min. Then, the partition was removed, and the behaviors were recorded until the pup escaped into the right chamber. Two persons who did not know the experimental conditions watched the video and examined the following two points: whether the rats left the right chamber and touched the boundary between the center and left chambers during the habituation, and whether the test rats touched the pups within 30 s after the partition removal.

#### 4RT

For the 4RT, we constructed a 60-cm square acrylic box partitioned into four rooms using black opaque acrylic boards. A rat pup of PND11 that was unknown to the test rats was placed in the corner of one of the four rooms, and immediately after, the test rat was placed in the center of the square box. The illuminance was 41 and 24 lux at the center of the apparatus and in the corners, respectively. The pups were only used once for the test. The rats’ behavior was recorded for 5 min by overhead video recording. Two persons who did not know the experimental conditions visually measured the time spent by the pups and the test rat in the same compartment. The measurements of the two persons were then averaged.

#### RarT

We created a transparent acrylic rectangular box (28 × 7 × 6 cm), with doors on the front and back (Fig. 3Ga). The doors could only be opened by pushing from the outside, not from the inside. The door was marked with vertical black lines so that the rats could easily recognize it. A male rat of the same age as the test rats was restrained in the box, which was fixed on the floor of the OF arena where the comatose rat in TT were fixed, and the same area (40 × 25 cm) as the Live zone in TT was designated as the Rescue zone. Then, a test rat was released from one of the corners of the OF arena, and the movement was recorded for 5 or 10 min using the video-tracking system and a video camera. A restrained rat was used for two tests involving the LR and Cont rats. The duration spent in the Rescue zone and the time of first entry into the zone were measured using the video-tracking system. Illuminance was the same as that in the OFT.

#### BT

An observation box (40 × 30 × 30 cm) for BT was constructed using acrylic boards (Fig. 4A). The rear and right sides were made with translucent gray acrylic boards, where an experimenter could observe the animals’ behavior. An opaque black acrylic board with a rough surface was used for the bottom of the box. The other parts were made of transparent acrylic boards. At the top, we placed a transparent acrylic lid with 4 cm-diameter holes in four corners to prevent animals from escaping. Using two removable partitions, we divided the apparatus into three rooms: the ICR, BL6, and Rat rooms. The center of the apparatus was illuminated at 47 lux. The partitions had holes with a diameter of 1 cm at approximately 4.5 cm intervals, 5 cm from the bottom, so that the animals could smell each other. A cosmetic brush that was attached to a 1cm-square acrylic rod (40 cm long) was used to stop any dangerous attacks between the animals that may cause injuries. Video cameras were set up on the front and left sides of the box to record the animal behaviors. The floor of the box was covered with a 2 cm-thick floor bedding. The test was conducted using a 11- to 13-week-old male ICR and a male BL6 younger by 1 week than ICR in addition to the test rat. First, a bullying pair of ICR and BL6 was placed in the box without partitions to confirm that the ICR attacked the BL6 within 1 min. Then, the mice were removed, and a test rat was put into the box, allowing them to move freely for 2 min to acclimate to the environment as well as the brush, which was moved slowly in front of the rat (Fig. 4B). Next, we inserted a central partition to separate the rat and mouse rooms and then placed the bullying pair in the mouse room side so that the rats could observe the bullying by the ICR for 3 min. If the ICR attacks did not reach 5 times, the time was extended until five attacks occurred. Thereafter, we separated the mice by inserting a partition between the ICR and BL6 rooms. The rat was left in this condition for 2 min to calm itself down from agitation caused by watching the attack. Finally, we removed all the partitions to allow the three animals to contact each other, and observed them for 3 min. During this time, violent contacts between the animals may occur, but could be stopped using the brush (fig. S9). In the experiments with the peaceful pairs were conducted after completing those with the bullying pairs, rat habituation was omitted. The partition between the mouse and rat rooms was then inserted to allow the rat to observe the interaction of the peaceful pair for 1 min. We measured the number of attacks by the ICR on the BL6 for 3 min and the number of contacts with the ICR or BL6 by the rats’ forepaws. After removing the rat from the box, we recorded the behavior of the two mice for 1 min. Three persons (an undergraduate, a laboratory staff, and a secretary) who were unfamiliar with the experimental conditions counted the number of contacts by the test rats with the ICR or BL6 and the number of ICR attacks for 3 min when the three animals could contact each other. They also examined whether the ICR attacked the BL6 within 1 min after the rat was removed.

### Statistical Analysis

Data are expressed as the mean ± SD. Statistical data were analyzed using two-tailed unpaired or paired *t*-test, χ^2^ test with Fisher’s exact test, one- or two-way analysis of variance (ANOVA) with Tukey’s post hoc test, or Spearman correlation analysis (the method used in each experiment is described in the figure legends). All analyses were conducted using Prism 8 (GraphPad Software, La Jolla, CA, USA). A *p*-value less than 0.05 was considered significant for all tests. The single, double, triple, and quadruple asterisks in the graphs indicate statistical significance at *p* < 0.05, 0.01, 0.001, and 0.0001, respectively.

## Acknowledgements

The authors thank Hirotomo Nakaoka and Chie Shiraishi of the Animal Center, Ehime University, for the maintenance of the facilities.

## Funding

This work was supported by the Japan Society for the Promotion of Science (JSPS) Grant-in-Aid for Challenging Research (Exploratory) 20K21465 to JT; Grant-in-Aid for Scientific Research (C) to TN (19K09436; 23K08383).

## Author contributions

Conceptualization: JT

Data curation: KM, YK, HY, JT Formal Analysis: KM, YK, TD, JT Funding acquisition: TN, JT

Investigation: KM, YK, TD, MEC, YN, RT, YW, HK, MK, JT

Methodology: KM, NH, JT Project administration: JT Resources: MEC, JT Supervision: JT Validation: KM, JT Visualization: KM, YK, JT Writing – original draft: JT

Writing – review & editing: TN, JT

## Competing interests

Authors declare that they have no competing interests.

## Data and materials availability

The majority data are available in the main text or the supplementary materials. Any other data will be provided upon request. In particular, we are pleased to send large-sized videos in response to requests through appropriate ways.

## Notes

### Competing Interest Statement

The authors have declared no competing interest.

