## Supplementary figures and images for "Wistar rats raised in an affectionate environment display lifesaving-like behaviors while distinguishing life from death"

### Fig. S1.png

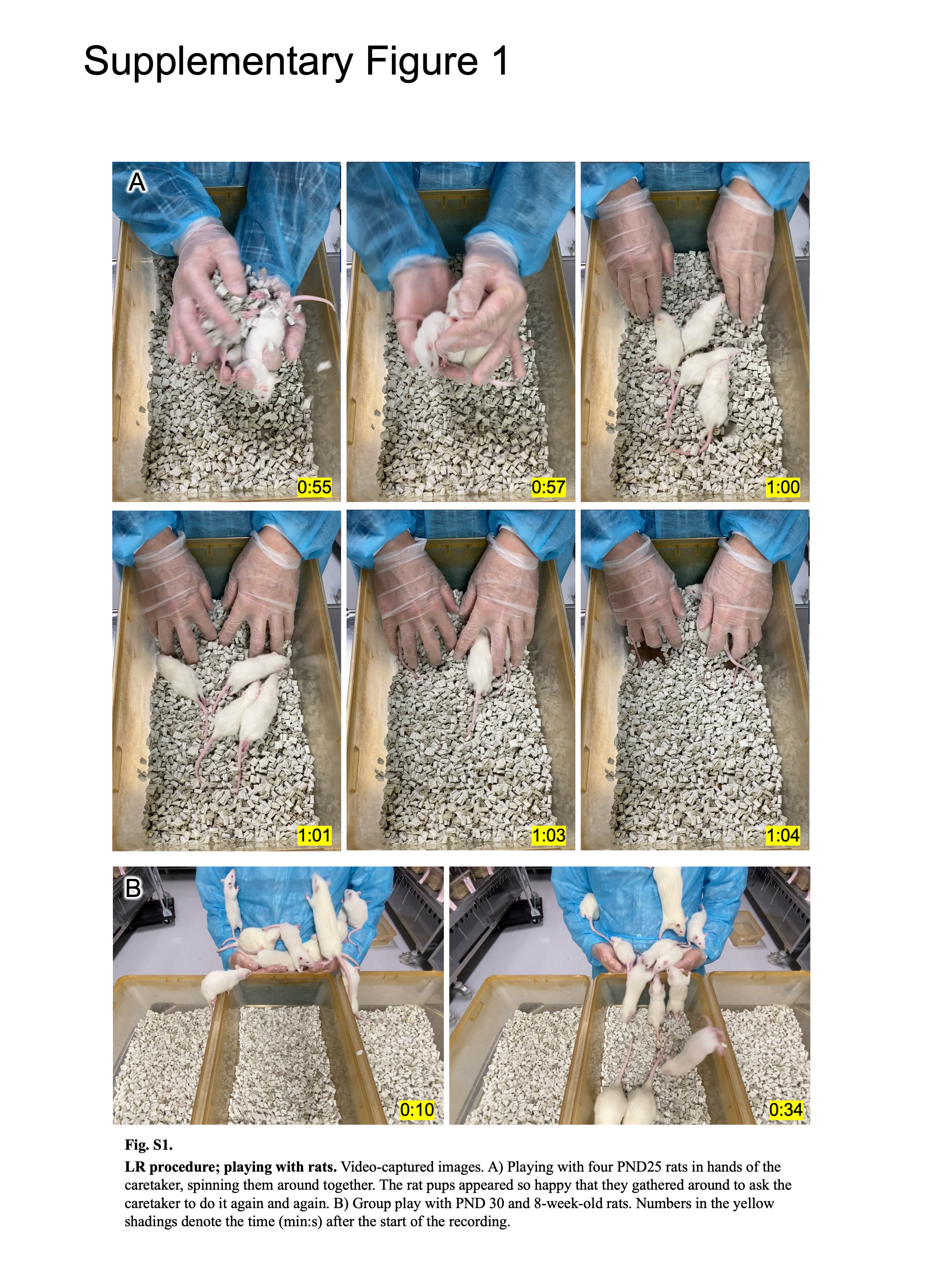

### Fig. S2.png

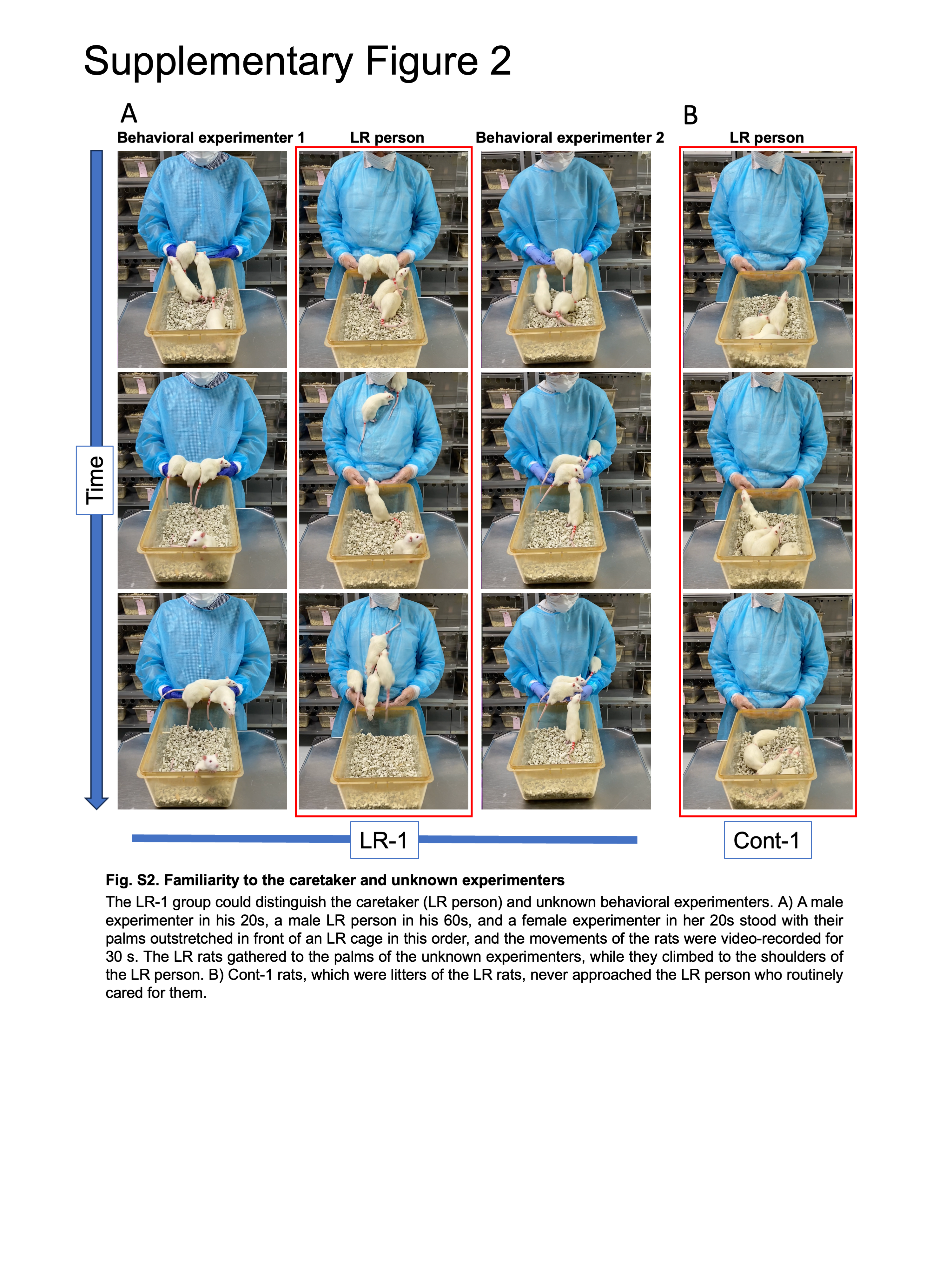

### Fig. S3.png

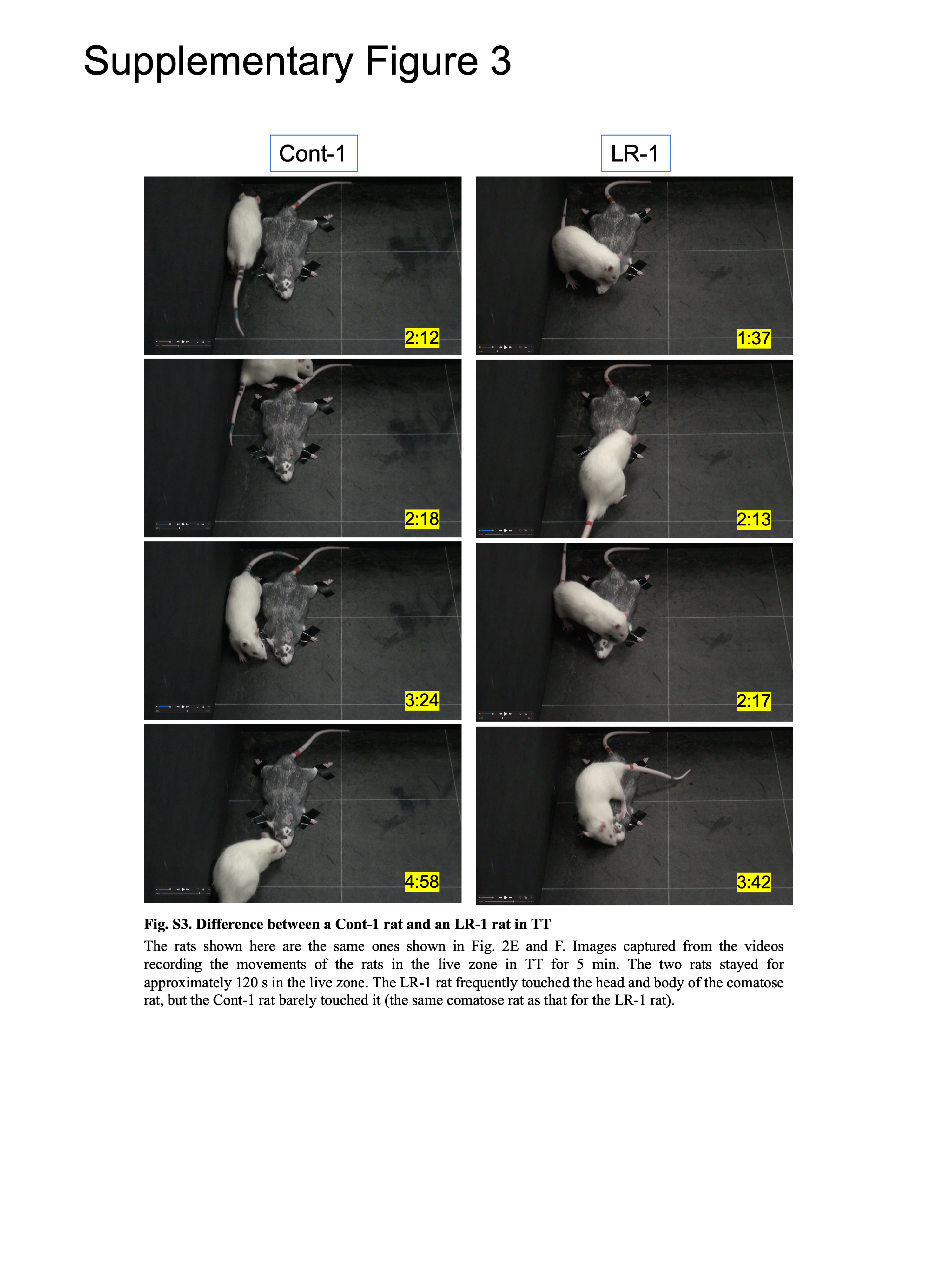

### Fig. S4.png

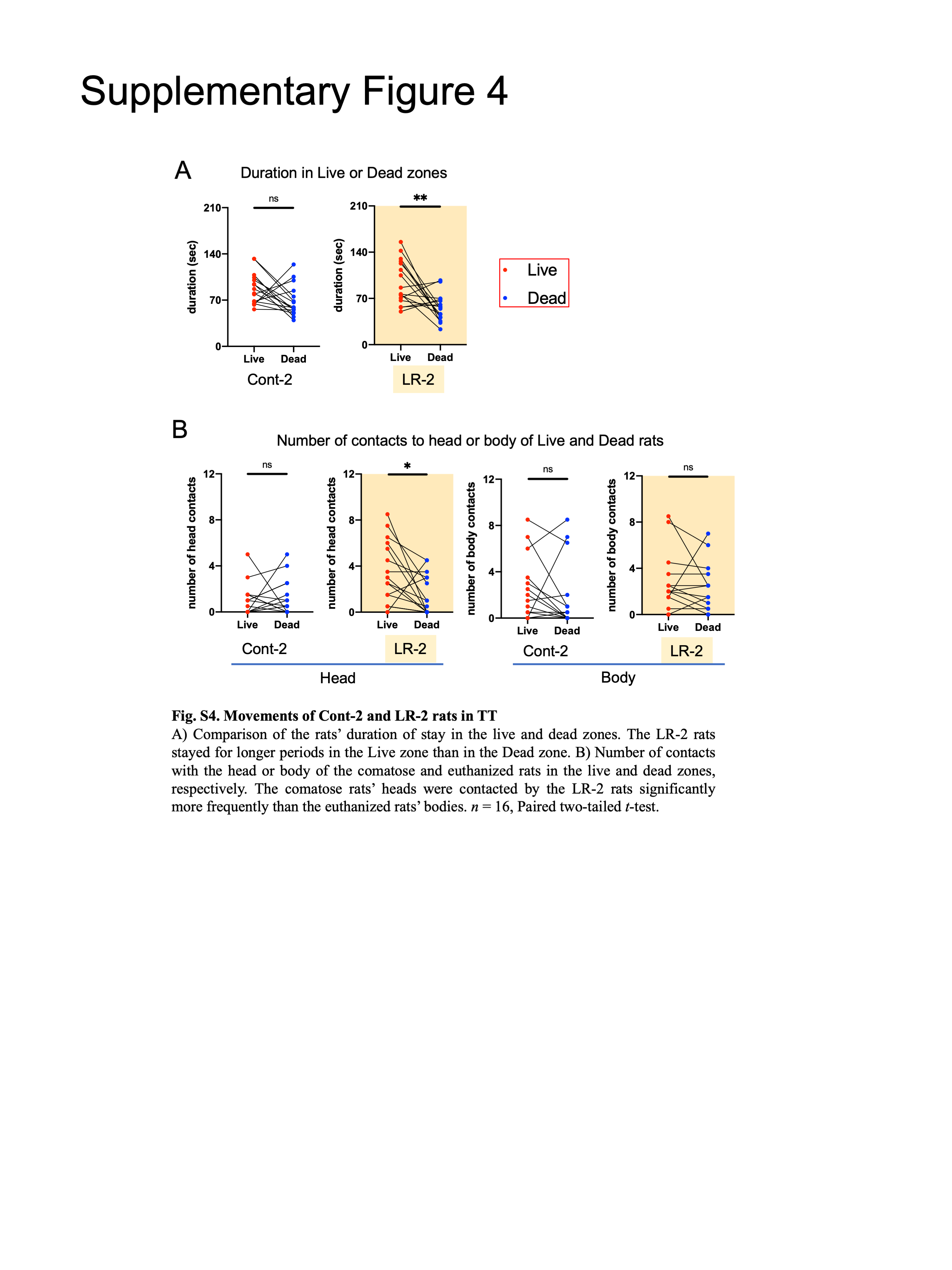

### Fig. S5.png

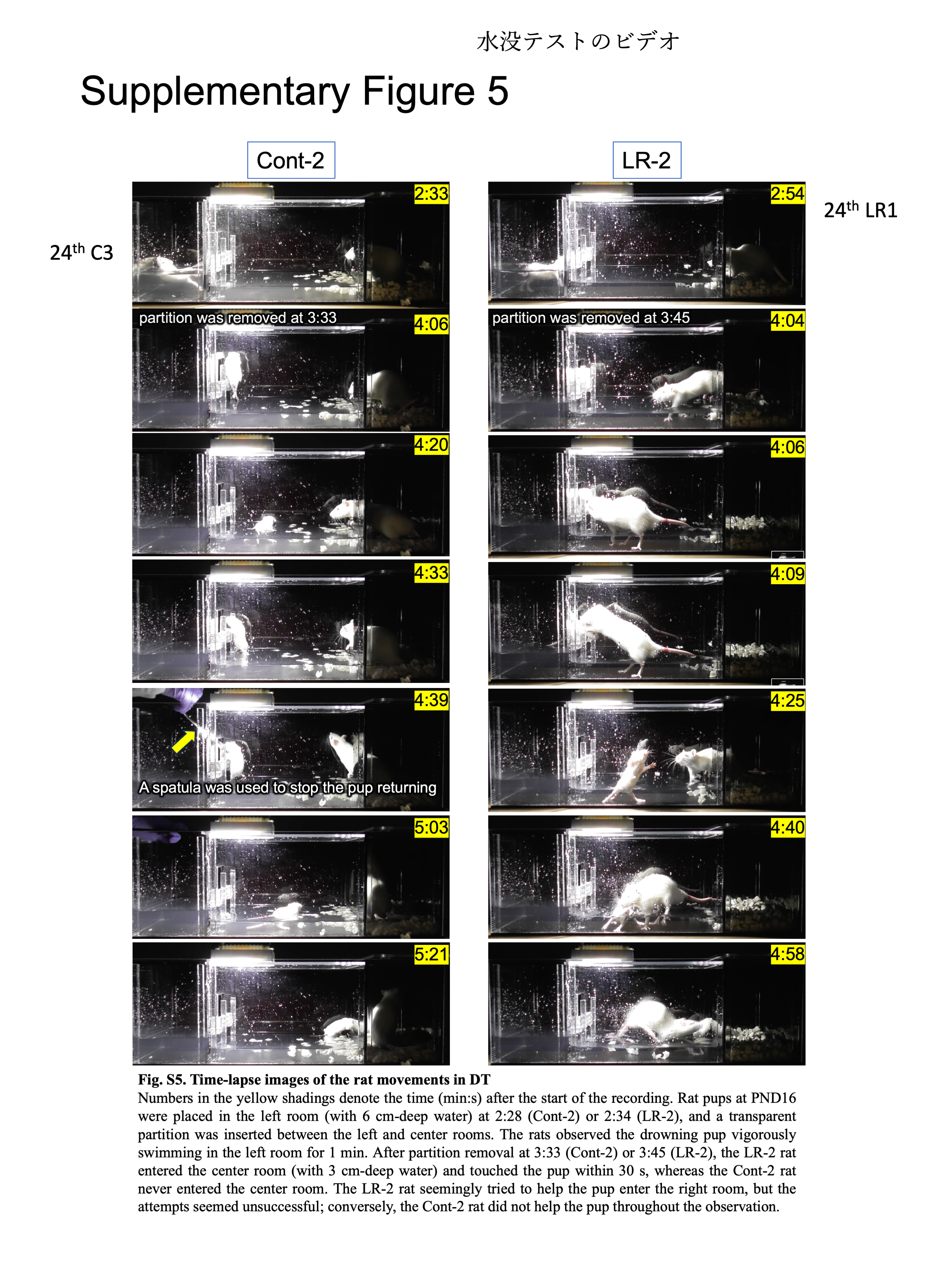

### Fig. S6.png

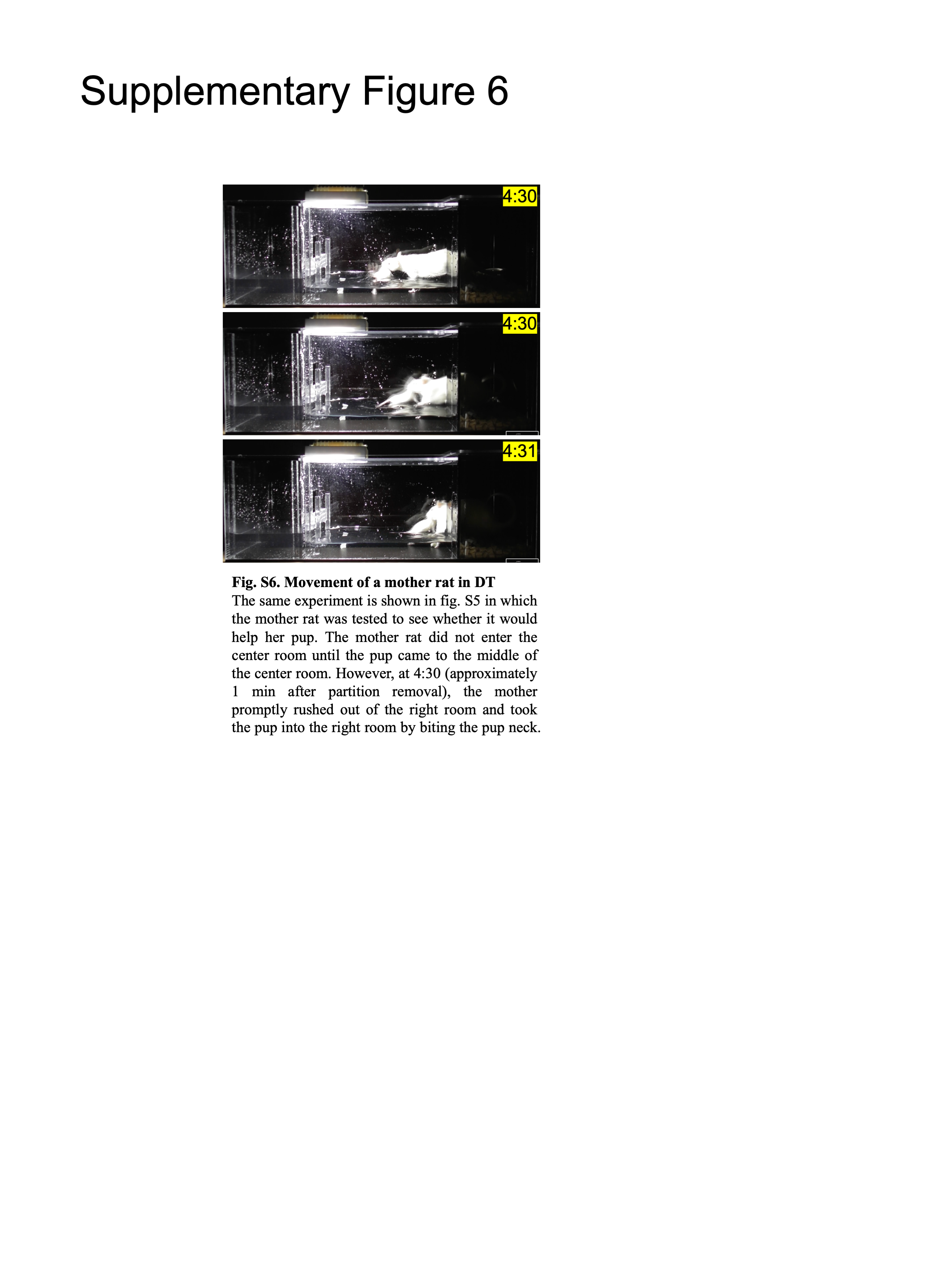

### Fig. S7.png

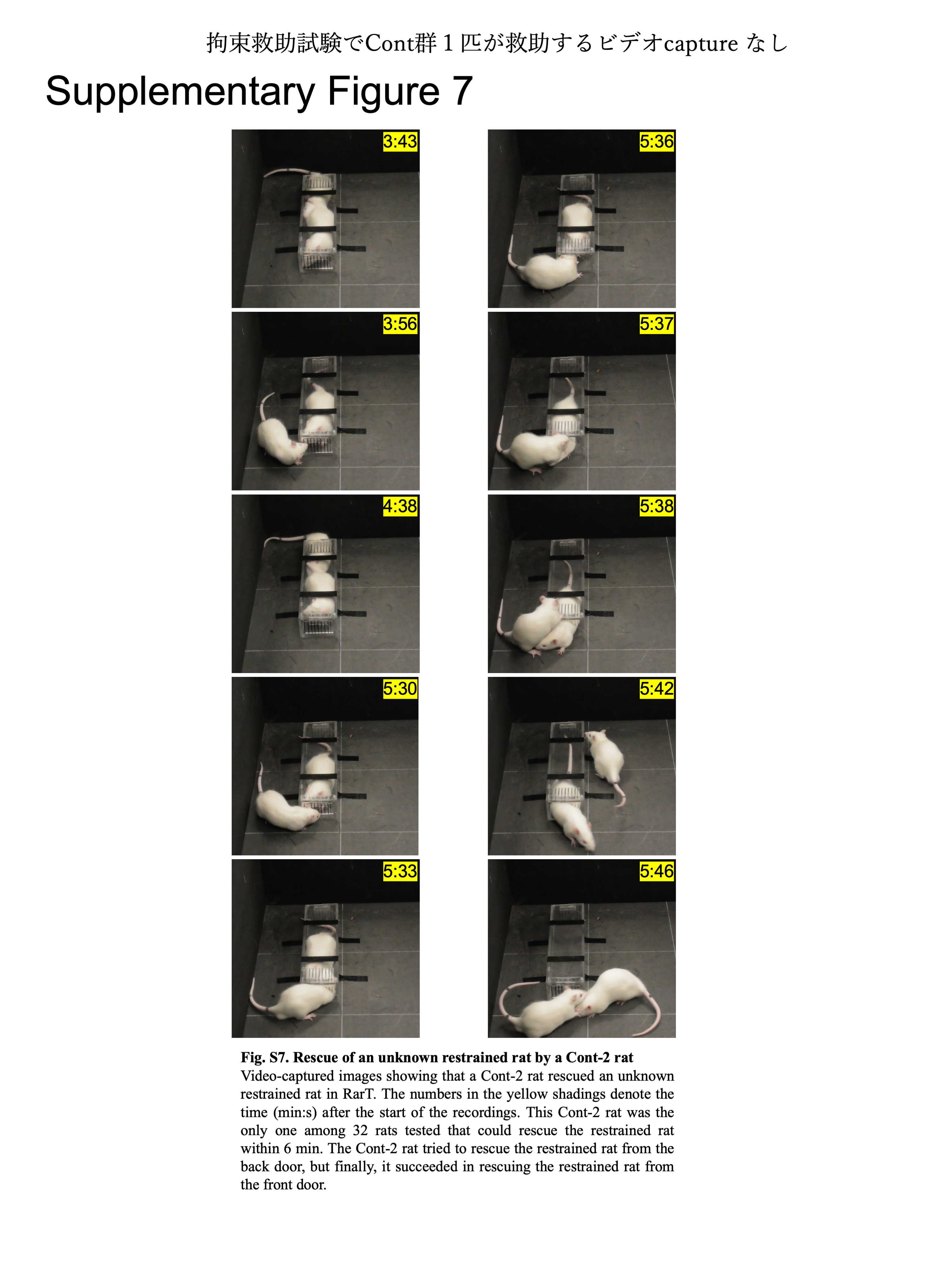

### Fig. S8.png

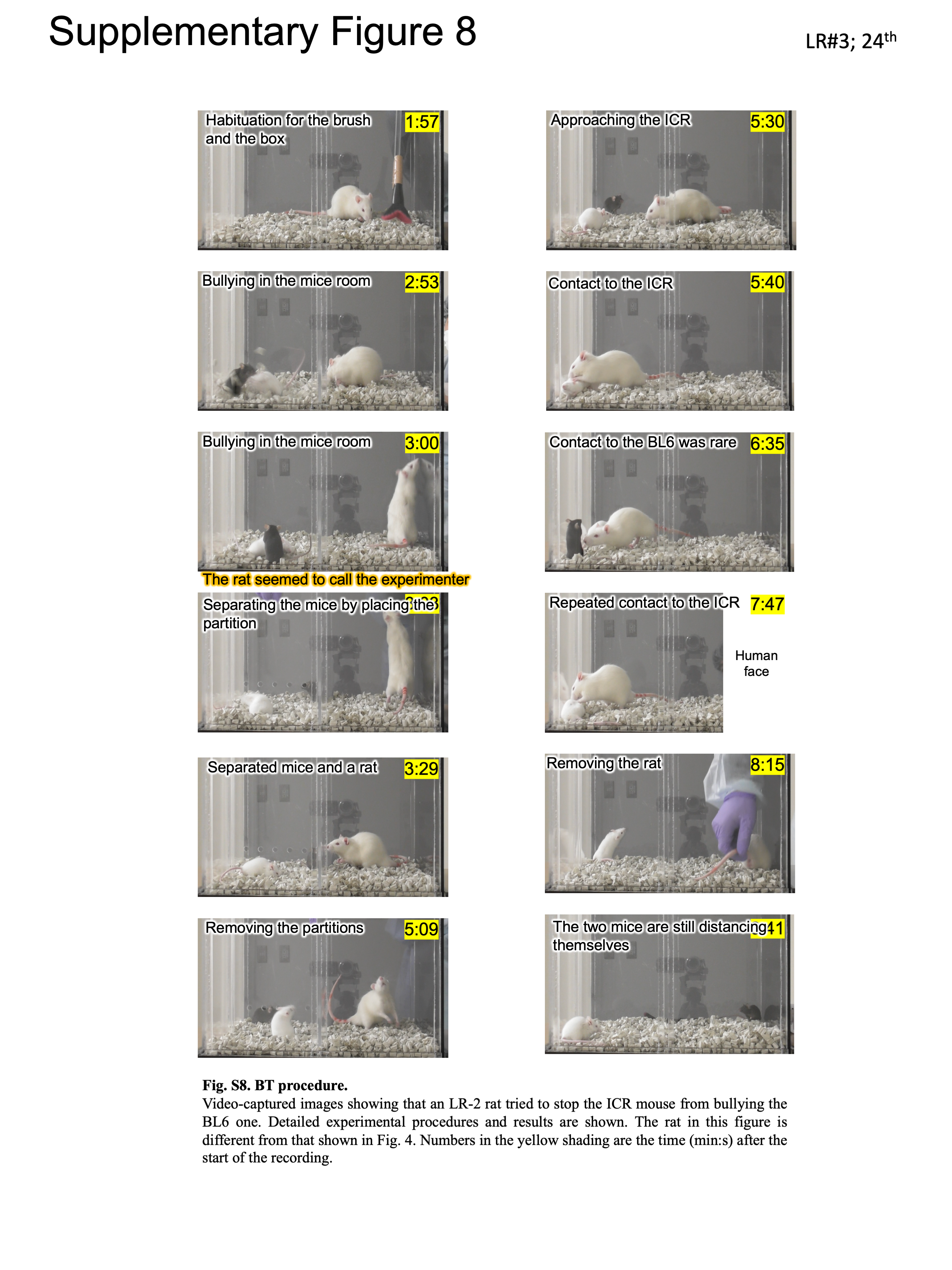

### Fig. S9.png

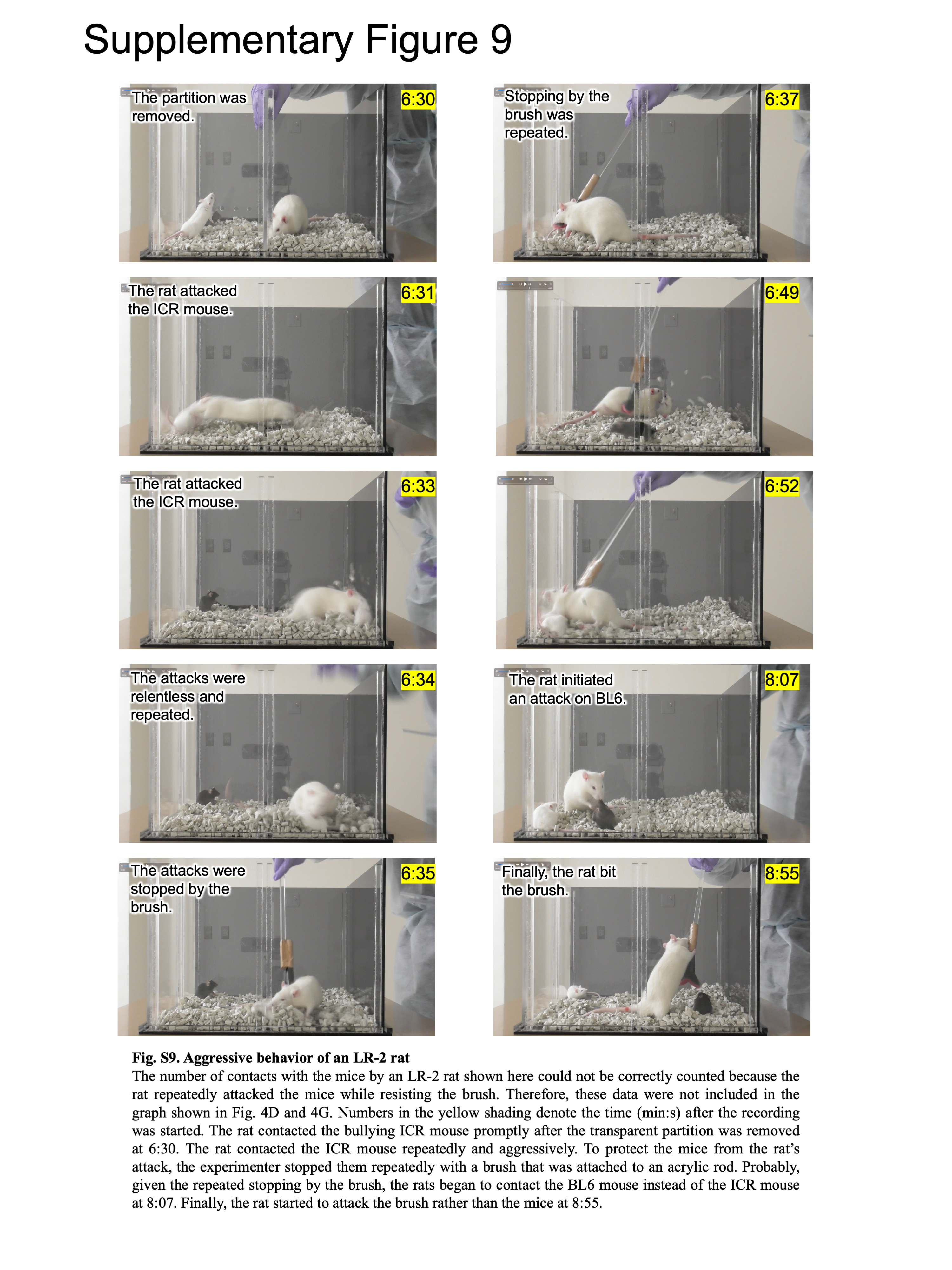
